# Recombineering in *C. elegans*: genome editing using *in vivo* assembly of linear DNAs

**DOI:** 10.1101/050815

**Authors:** Alexandre Paix, Helen Schmidt, Geraldine Seydoux

**Author notes:** Corresponding authors: Alexandre Paix, Geraldine Seydoux.

## Abstract

Recombineering, the use of endogenous homologous recombination systems to recombine DNA *in vivo*, is a commonly used technique for genome editing in microbes. Recombineering has not yet been developed for animals, where non-homology-based mechanisms have been thought to dominate DNA repair. Here, we demonstrate that homology-dependent repair (HDR) is robust in *C. elegans* using linear templates with short homologies (~35 bases). Templates with homology to only one side of a double-strand break initiate repair efficiently, and short overlaps between templates support template switching. We demonstrate the use of single-stranded, bridging oligonucleotides (ssODNs) to target PCR fragments precisely to DSBs induced by CRISPR/Cas9 on chromosomes. Based on these findings, we develop recombineering strategies for genome editing that expand the utility of ssODNs and eliminate *in vitro* cloning steps for template construction. We apply these methods to the generation of GFP knock-in alleles and gene replacements without co-integrated markers. We conclude that, like microbes, metazoans possess robust homology-dependent repair mechanisms that can be harnessed for recombineering and genome editing.

## INTRODUCTION

Recombineering (recombination-mediated genetic engineering) is a molecular engineering method used to manipulate the genomes of bacteria and yeast (1). Recombineering relies on the robust recombination machinery present in these organisms that allow exogenous DNAs to recombine with each other and with chromosomal loci with only minimal homology requirements. Recombineering methods are not available yet for animal cells. For genetic engineering in animal models, synthetic DNAs are assembled first typically in *E. coli* and are transferred in a second step into the animal (2–4).

CRISPR-Cas9 technology has simplified methods to introduce exogenous DNA into eukaryotic genomes (5). The RNA-guided endonuclease Cas9 creates a double-strand break (DSB) at a precise site in the genome that is repaired using the cell’s DNA repair machinery. If an exogenous DNA with homology to the cleavage site (“homology arms”) is provided at the same time, the cell’s repair machinery will use that DNA as a template to repair the DSB (homology-dependent repair or HDR). As a result of the repair process, sequences in the template between the homology arms will be inserted in the genome. Two types of repair templates have been used most commonly in animals: single-stranded oligonucleotides (ssODNs) with short (<60 nucleotides) homology arms and plasmids with longer (>500 bases) homology arms. ssODNs can be synthesized chemically, but can only be used for small (<130 bases) edits. Plasmid templates can accommodate larger edits (such as GFP) but must be assembled by the user. In contrast in microbes, it is possible to use recombineering to assemble templates *in vivo* from linear DNAs with short homologies. Combined with CRISPR/Cas9 technology, recombineering has been used to build efficient pipelines for the generation of complex genomic edits without selection (6). Methods that would allow recombineering in animal cells would simplify repair template construction and thus expand the possibilities offered by Cas9-assisted genome editing.

Studies in budding yeast have described a gene conversion mechanism for the repair of DSBs called synthesis-dependent strand annealing (SDSA), where DSBs are sealed using sequence copied from an homologous template (7). SDSA begins with resection of DNA ends in the 5’ to 3’ direction to expose single-stranded DNA on both sides of the DSB. The single-stranded ends pair with homologous sequences on another chromosome or an exogenously supplied donor DNA. After pairing, the invading strand is extended at its 3’ end by DNA synthesis using the homologous sequence as template. When sequences complementary to the other side of the break are copied, the newly synthesized strand withdraws and anneals with the other resected end on the chromosome. Synthesis of the other strand and ligation seal the break (8). In cases where a second donor template is also present, the newly synthesized strand can anneal to the second template and continue DNA synthesis (“template switching”), effectively stitching the two donor template sequences together (9). Here, we present evidence that a similar gene conversion mechanism with robust template switching operates in *C. elegans* during repair of Cas9-induced DSBs. We identify the homology requirements needed for linear DNAs to recombine with each other and with Cas9-induced DSBs on chromosomes. We build on these findings to develop recombineering methods for genome editing in *C. elegans*.

## MATERIALS AND METHODS

### Reagents and sequencing

Recombinant His-tagged Cas9::SV40 was purified following protocols in (10). crRNAs and tracrRNA were obtained from Dharmacon and reconstituted in 5mM Tris pH7.5 at 8μg/μl and 4μg/μl respectively. ssODNs were obtained from IDT (salt free purification). PCR amplicons were generated as described in (11). For several experiments (Supplementary Table S1), we selected representative edits for sequencing: the edits were made homozygous before PCR amplification and sequencing of the entire insert including both junctions (Supplementary Table S1). Sequences of crRNAs, ssODNs, PCR primers and inserts are shown in Supplementary Tables S2/S3/S4, and plasmids and strains are listed in Supplementary Table S5.

### Editing protocol

Editing experiments were performed following methods described in (10) using *in vitro* assembled Cas9 ribonucleoprotein complexes and the co-conversion method to isolate edits (12). Co-conversion uses co-editing of a marker locus (*dpy-10*) to identify animals derived from germ cells that have received Cas9 and the repair templates, reducing possible experimental noise due to variations in injection quality from animal to animal (10,12). We used a ~1/3 ratio of *dpy-10/locus of interest* crRNAs to maximize the recovery of desired edits among worms edited at the *dpy-10* locus (10). Injection mixes contained 15.5 μM Cas9 protein, 42 μM tracrRNA, 11.8 μM *dpy-10* crRNA, 0.4 μM *dpy-10* repair ssODN, 29.6 μM locus of interest crRNA(s) and varying concentrations of repair templates (0.1-0.5 μM; Supplementary Table S1). For *gtbp-1* replacement (Figure 4I), both 5’ and 3’end crRNAs were used at 22.2 μM each and the tracrRNA concentration was increased to 56.2 μM. Final buffer concentrations in injection mixes were 150mM KCl, 20mM HEPES, 1.6mM Tris, 5% Glycerol, pH7.5-8, except for Figure 2E/F/G and Figure 4A where KCl was at 200mM and for Figure 4I where Tris was at 2.1mM. Injection mixes were assembled by mixing the components in the following order: Cas9 protein, KCl, HEPES pH7.5, crRNAs, TracrRNA, ssODNs, H2O and finally PCR fragments if used.

**Figure 4:**
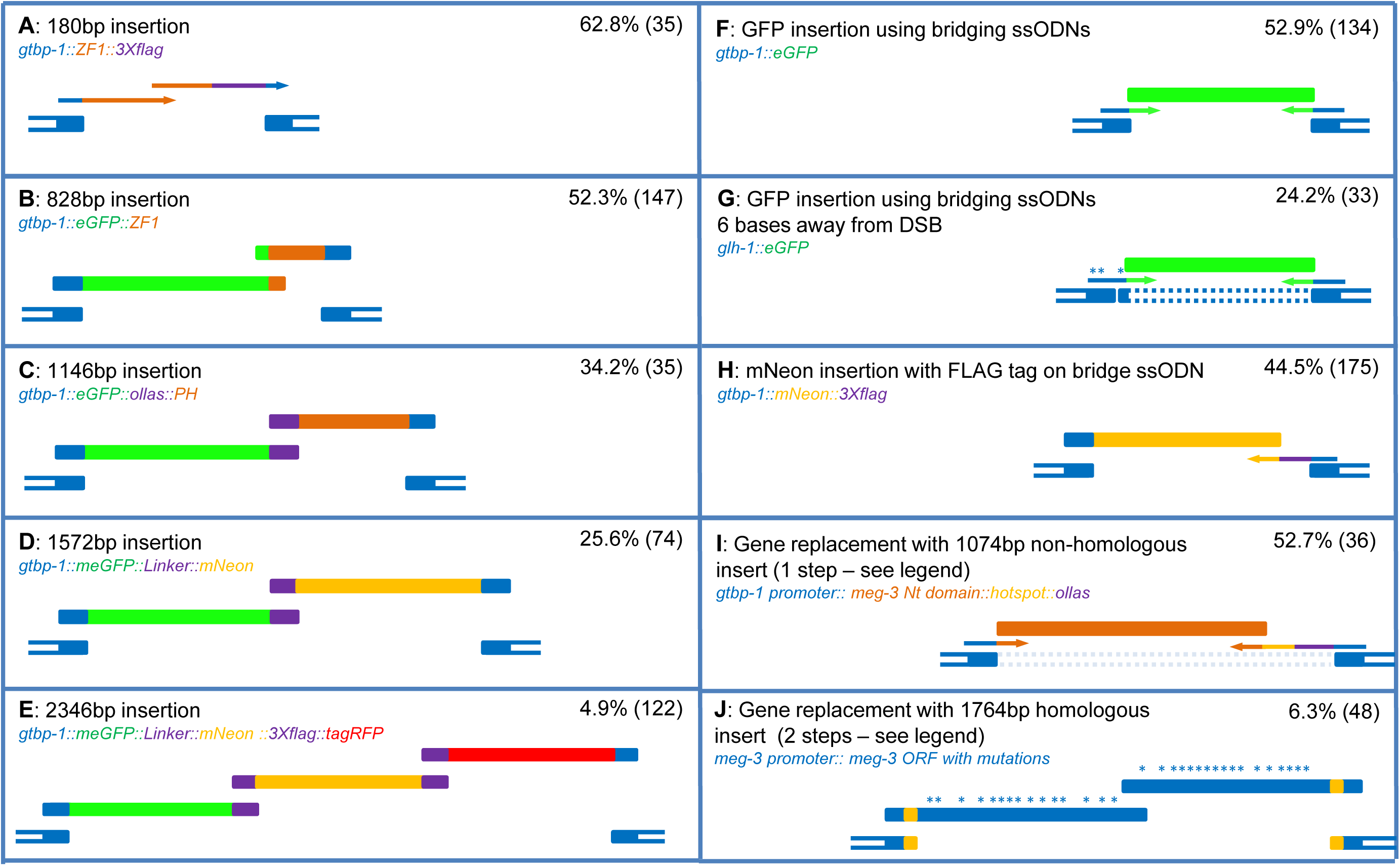
Recombineering applications. Schematics (not to scale) show the targeted DSB (bottom) and repair templates (top): dark blue are genomic sequences, and other colors are as indicated in each panel. Bridging ssODNs are represented with one line with arrow at 3’ end. All overlaps are ~35 bases .% edition correspond to the number of worms edited at the locus depicted among *dpy-10* edited worms. In panel G, dotted double lines connect contiguous sequences in the *glh-1* locus and stars represent recoded sequence to avoid recutting by Cas9. In panel I, two Cas9-induced DSBs were used to delete the *gtbp-1* ORF (light colored dotted line) and replace it with non-homologous sequence (brown) in one step. In panel J, this gene replacement was performed in two steps. In the first step (not shown), the *meg-3* ORF was deleted using two DSBs and repaired with a bridging ssODN containing a new Cas9 recognition site (orange). In the second step (shown), the mutated *meg-3* ORF was reinserted at the new Cas9 site using two PCR fragments. PCR fragments were injected at 0.44-0.48 pmol/μl except for panel D (0.34 pmol/μl), panel E (0.18-0.20 pmol/μl), panel I (0.31 pmol/μl) and panel J (0.32 pmol/μl). Additional information including sequencing results can be found in Supplementary Table S1.

**Figure 2:**
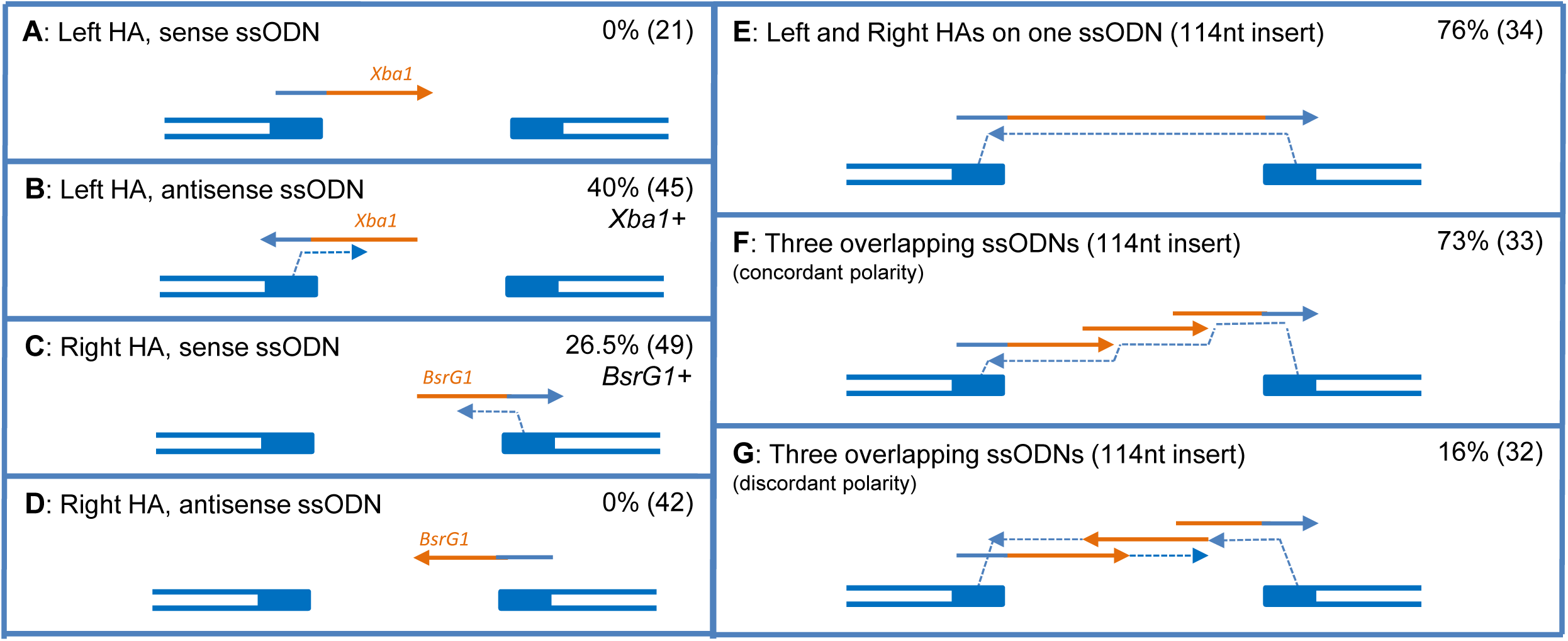
Usage of homology arms on ssODNs and evidence for template switching. Double lines represent genomic sequences and single lines are repair ssODNs (arrows at 3’ end). Blue are genomic sequence or sequences on ssODNs homologous to genomic sequences, brown are unrelated sequences. Fine stippled lines and arrows show sites of strand invasion and replication inferred from the type of edits recovered Percentages represent the % of worms edited as shown among *dpy-10* edits (n= number of *dpy-10-*edited worms screened). Edits were identified by PCR genotyping of F2 cohorts derived from cloned single F1 worms. Additional information can be found in Supplementary Figures S2/S3 and Supplementary Table S1.

Each injection mix was injected in the oogenic gonad of ~20 isogenic and synchronized young adult hermaphrodites (wild-type N2 or *meg-3* deletion in Figure 4J). The injected mothers were cloned to individual plates 24 hours after injection. 5-6 days later, broods with the highest numbers of *dpy-10* edits were identified (“jackpot broods”). This step selects for broods derived from hermaphrodites that were injected successfully. For each experiment, *dpy-10*-edited progenies from at least three independent jackpot broods were screened for edits at the locus of interest. GFP+ edits were identified by direct inspection of adult F1 animals for GFP expression in the germline. All other edits were identified by PCR genotyping of F2 cohorts derived from cloned F1s. All edits reported were germline, heritable edits. The majority of edits were recovered in the heterozygous state in F1 progenies, but we also obtained a minority of homozygously edited F1s. These observations show that, as expected, edits are created primarily shortly after injection in the oogenic germline (the site of injection). Occasionally, however, edits are also created on paternal chromosomes, presumably in zygotes shortly after fertilization since all edits were germline edits (inherited by next generation). These observations confirm that homology-dependent repair also occurs in zygotes, using the donor templates or the previously edited maternal allele, as we have observed previously (10).

### Calculation of editing efficiency

Editing efficiency was calculated as the percentage of *dpy-10*-edited F1 progenies that were also edited at the *gtbp-1* locus or other locus of interest. This method normalizes the edit efficiency at the locus of interest against the edit efficiency at the marker *dpy-10* locus to minimize the effect of possible variation in Cas9 activity or delivery between experiments. Differences in editing efficiencies between different templates reflect differences in the templates abilities to support HDR. We have confirmed that our methods yield reproducible editing frequencies when comparing the same injection mix injected on different days (for example, see Supplementary Table S1 for replicates of experiments summarized in Figure 1A and 2G).

**Figure 1:**
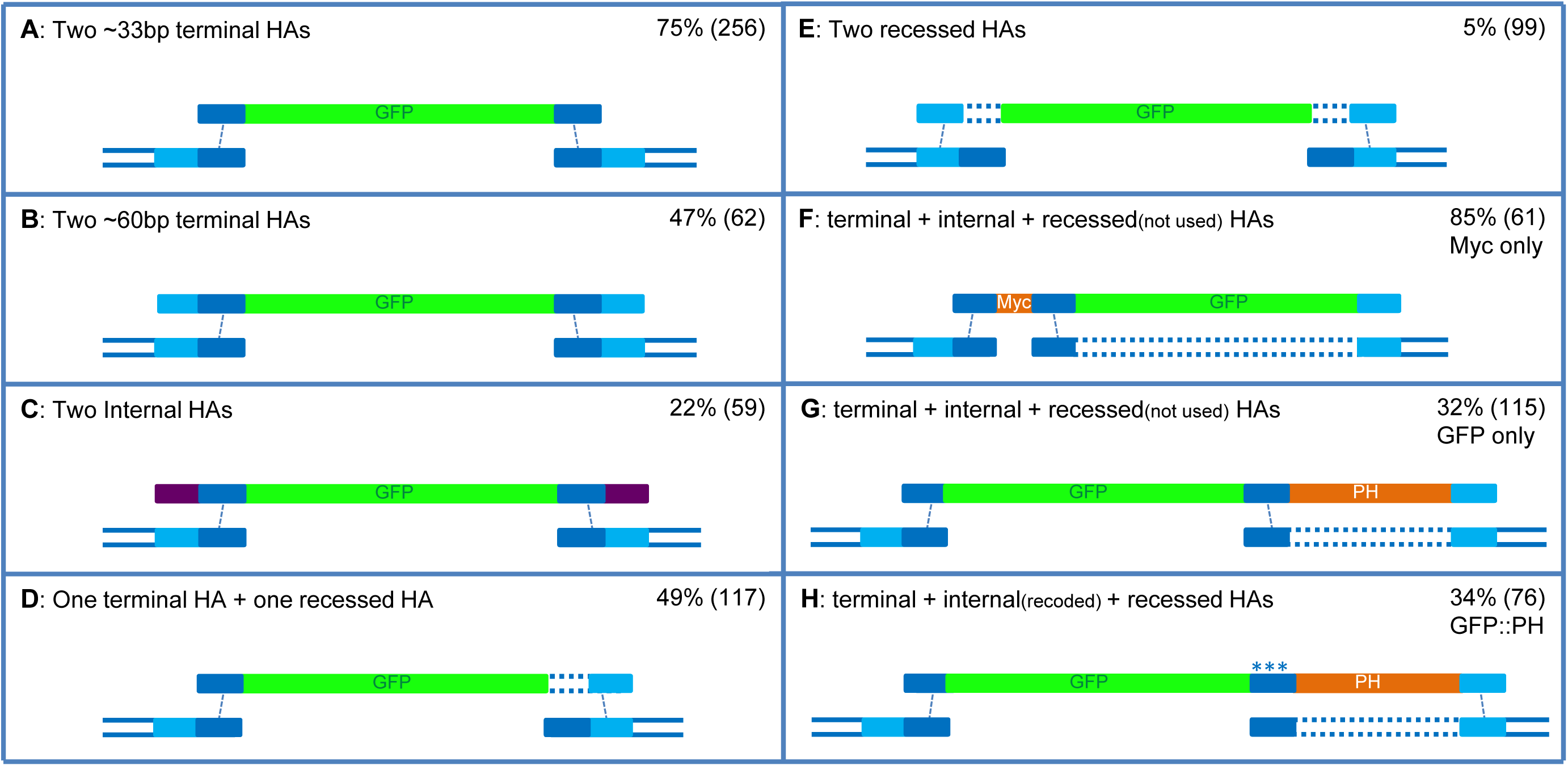
Usage of homology arms on PCR templates. In this and all subsequent figures, each schematic shows the repair templates (top) and the DSB at the genomic locus (bottom) aligned by homology. Percentages are the percent of worms edited as shown among *dpy-10* edits (n= number of *dpy-10-*edited worms screened). Blue are genomic sequence or sequences on templates homologous to genomic sequences, all other colors are unrelated sequences. Dark and light blue: proximal/distal homology arms (HAs) relative to DSB, asterisks: recoded sequence, dotted double lines connect contiguous sequences in the genome/template that are separated in the diagram to line up with homologous sequences in the template/genome. Fine stippled lines show sites of strand invasion inferred from the type of edits recovered. All homologies/overlaps are ~35 bases long (see Supplementary Table S1 for details).

The maximal editing frequency we observed at the *gtbp-1* locus (among *dpy-10* edits) was 85% (Figure 1F). Because we do not know the *in vivo* efficiency of the *gtbp-1* crRNA, we do not know whether this percentage reflects the maximal efficiency of HDR or the maximal efficiency of introducing DSBs at the *gtbp-1* locus, in which case HDR efficiency at DSBs could be even higher.

## RESULTS

### HDR favors templates with terminal homology arms that correspond to sequences directly flanking the DSB

Previously we demonstrated that PCR amplicons with short (~35-base) homology arms can be used as repair templates for Cas9-induced DSBs on chromosomes (11). To determine the optimal design of homology arms, we compared the performance of donor templates designed to insert a promoterless GFP at a single Cas9-induced DSB near the C-terminus of the *gtbp-1* locus. Only in frame, precise insertions will give rise to GFP+ edits (10). To maximize the generation of DSBs, we injected Cas9, and associated crRNAs and tracrRNAs, as ribonucleoprotein complexes into the germline of adult hermaphrodites and used co-conversion of the marker locus *dpy-10* to identify progeny derived from Cas9+ germ cells (12) (10). Editing efficiency was calculated as the % of *gtbp-1* edits among progeny edited at the *dpy-10* locus. This method normalizes edit efficiency across experiments (Material and Methods) and provides a metric to compare the competence of different templates to engage in HDR (Materials and Methods). We obtained the highest percentage of GFP+ edits with a PCR fragment where GFP was flanked by two ~35-base homology arms corresponding to sequences directly surrounding the DSB (two “terminal” homology arms: 75% GFP+ edits, Figure 1A). As we observed previously using a different method to isolate edits (11), longer homology arms reduced editing efficiency (60 bases: 47%, 100 bases: 33%, Figure 1B and Supplementary Table S1). Capping both homology arms with 33 bases of non-homologous sequence (to create “internal” homology arms) also decreased efficiency (22%, Figure 1C and Supplementary Table S1). Templates with homology arms corresponding to sequences ~30 bases away from the cut (“recessed” arms) also performed poorly (Figures 1D and 1E). Loss of efficiency was most pronounced when both arms were recessed (5%, Figure 1E). These observations confirm that the optimal homology arms are: short (~35-bases), located at the termini of the PCR fragment, and corresponding to sequences directly flanking the DSB.

Previously, we observed that templates with one homology arm that spans the DSB and one recessed arm designed to insert GFP at a distance from DSB do not yield GFP+ edits efficiently (10,11). To monitor the utilization of such templates, we placed a 30-base insert (Myc) in the arm spanning the DSB and screened for edits by PCR. We obtained 85% edits; all contained the Myc insert and none contained GFP (Figure 1F and Supplementary Figure S1). Similarly, we found that a template containing GFP flanked by two homology arms followed by a membrane localization (PH) domain and a recessed arm yielded only edits that expressed cytoplasmic GFP (Figure 1G). In contrast, the same template with mutations in the internal homology arm between GFP and the membrane localization domain yielded edits that all expressed membrane-bound GFP, confirming that the recessed arm can be used (Figure 1H and Supplementary Figure S1). These observations indicate that homology arms proximal to the DSB are preferred over recessed homologies, even when located internally to the template. In all cases, templates with at least one terminal homology arm performed better (Figure 1A/B/D/F/G/H) than templates with only internal homologies or only recessed arms (Figure 1C and 1E), suggesting that at least one terminal homology is required to initiate high efficiency HDR.

### Homology to one side of the break is sufficient to initiate repair and template switching can link overlapping templates

To investigate further the requirements for HDR, we used single-stranded oligonucleotides (ssODNs) as templates. Unlike PCR fragments, which are double-stranded and therefore can pair with both sides of the resected DSB, ssODNs are predicted to anneal to only one side of a resected DSB, either the right or left side depending on the polarity of the ssODN. To test this prediction, we compared ssODNs of opposite polarity and containing only one homology arm, corresponding to either the right or left side of the *gtbp-1* DSB (Figure 2). To monitor usage of the ssODNs, we looked for edits that incorporated a restriction site (RE) contained on the ssODN. As expected for templated-repair on one side of the DSB and non-templated repair on the other side of the DSB, we obtained edits of various sizes (Supplementary Figure 2). Edits containing the restriction site appeared only when using ssODNs with the correct polarity to pair with the resected end (Figure 2A-D, Supplementary Figure 2). Sequencing of 5 edits confirmed that HDR occurred on the side of the homology arm and NHEJ occurred on the other side (data not shown). These results are consistent with a repair process involving DNA synthesis templated by the ssODN (Supplementary Figure 2). These observations also demonstrate that templates with homology to only one side of the DSB can initiate HDR.

During SDSA in yeast, the newly synthesized strand can withdraw from one template and resume DNA synthesis on a second template (template switching; (9)). In principle, template switching could be used to link overlapping ssODNs. To test this hypothesis, we used three ssODNs to repair the single DSB at *gtbp-1*. The external ssODNs each had a 35-base homology arm corresponding to the right or left side of the DSB, and a 35-base overlap with the internal ssODN. We obtained 73% edits, a frequency similar to that obtained using a single ssODN to create the same 114-base insertion (76%, Figure 2E-F). Interestingly, we obtained only 16% full-size edits when the polarity of the internal ssODN was reversed (Figure 2G). In that configuration, we also obtained 12.5% partial edits, sequencing revealed that all contained sequences from the right most ssODN and none contained sequences unique to the left most ssODN (Supplementary Figure S3). These results are consistent with a repair mechanism initiated by ssODN pairing on the right side of the DSB followed by DNA synthesis and template switching that is most efficient when all ssODNs have the same polarity. We conclude that template switching can be used to link overlapping non-complementary ssODNs.

### Recombineering using overlapping PCR fragments and ssODNs

To determine whether templates can also be assembled from overlapping PCR fragments, we tested whether GFP could assembled from two ~400bp fragments with a ~35-base overlap in the middle. Each fragment also had a single ~35-base terminal homology arm corresponding to the left or right side of the DSB in *gtbp-1*. We obtained 57% GFP+ edits (Figure 3A) compared to 75% edits when using a single continuous GFP template inserted at the same site (Figure 1A).

**Figure 3:**
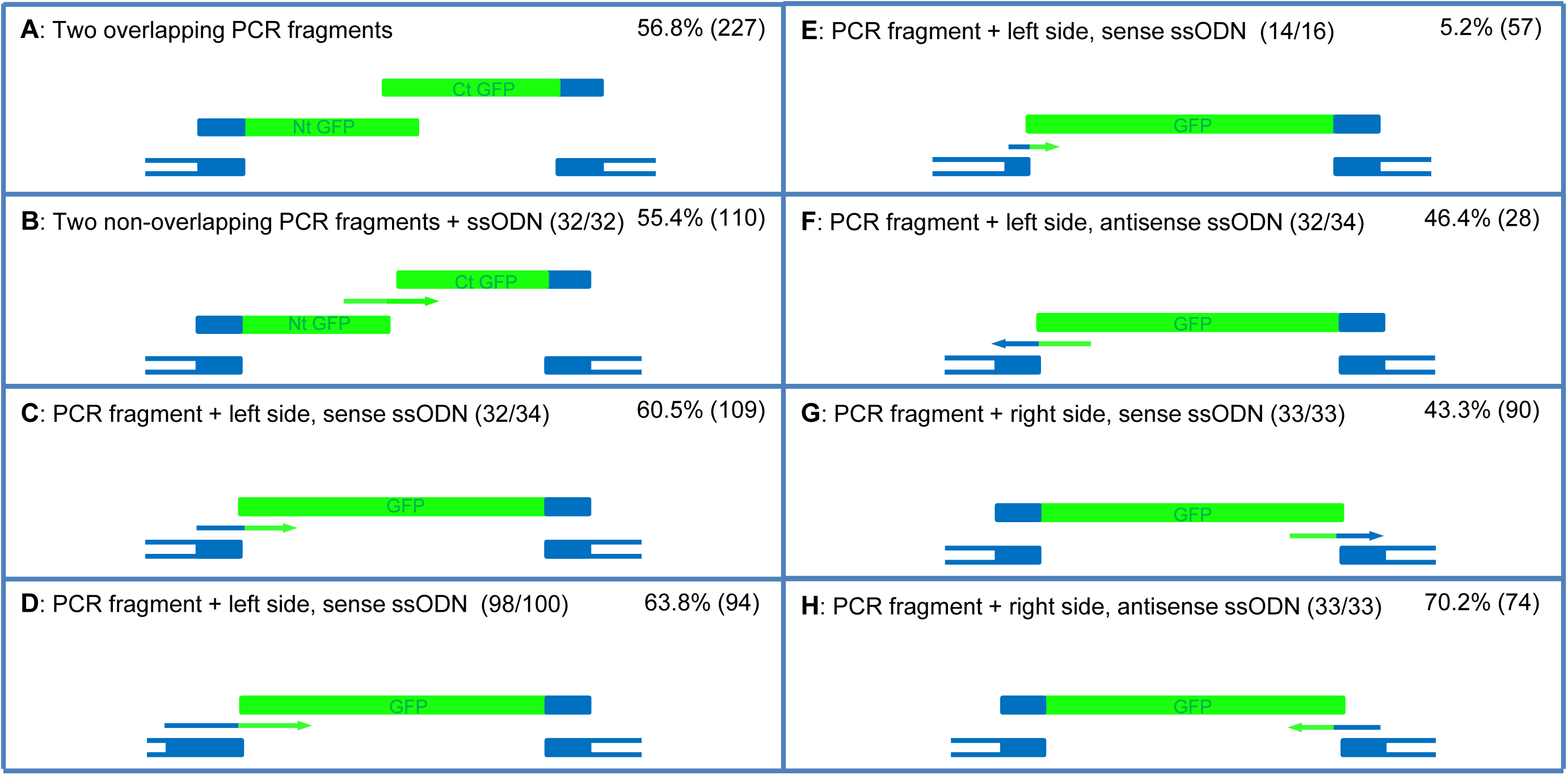
Recombineering using PCR fragments and ssODNs. Schematics (not to scale) show the targeted DSB and repair templates for each experiment: dark blue is *gtbp-1* genome sequence and green is GFP. Bridging ssODNs are represented with one line with arrow at 3’ end. Size of homology arms on ssODNs are indicated in parentheses. All other homology arms are ~35 bases. % edition correspond to the number of GFP+ worms among *dpy-10*-edited F1s. A GFP PCR fragment with no homology arms and no bridging ssODN gave 0% edits (Supplementary Table S1). See Supplementary Figure S4 for sequences of ssODNs and predicted pairing patterns.

Next we tested whether an ssODN could be used to link two fragments. We used two half-GFP PCR fragments that abutted but did not overlap and provided the overlap on a “bridge” ssODN that fused 2 x ~35 bases on either side of the GFP split. We obtained 55% GFP+ edits (Figure 3B), an efficiency similar to that obtained with the overlapping PCR fragments (Figure 3A). This result suggests that homologies to promote template switching between PCR fragments can be provided just as efficiently *in trans* (in ssODNs) as in *cis* (at the ends of PCR fragments).

Next we tested whether ssODNs could also be used to target a PCR fragment to a DSB on the chromosome. We used a PCR template with only one homology arm corresponding to the right side of the DSB in *gtbp-1* and used a bridge ssODN to provide homology to the left side of the DSB. The bridge ssODN contained 34 nucleotides homologous to GFP and 32 nucleotides homologous to the DSB. We obtained 60% GFP+ edits (Figure 3C), compared to 75% edits when using a PCR template with two homology arms and no bridge ssODN (Figure 1A). A control PCR template with no homology arms and no bridge ssODNs gave 0% edits (Supplementary Table S1). A longer ssODN with 2 x ~100 nucleotides homology arms did not perform significantly better than the ssODN with ~35 homology arms (compare Figure 3C and D), and a shorter ssODN (2 x ~15 nucleotides) performed poorly (5%) (Figure 3E). An ssODN of the opposite polarity gave 46% edits (Figure 4F). We also compared two ssODNs of opposite polarity for bridging to the other side of the *gtbp-1* DSB and again observed a slight preference for the ssODN that could pair with the resected template (Figure 4 G-H and Supplementary Figure S4). We conclude that ssODNs can be used to provide homology arms to target a PCR template to a DSB.

### Recombineering applications

The findings above demonstrate the feasibility of assembling templates *in vivo* from combinations of ssODNs and PCR fragments (recombineering). We have tested the utility of recombineering in the context of two common editing tasks: targeted insertions and gene replacements.

#### Small insertions using overlapping ssODNs

The current maximum size of commercially available ssODNs (200 bases) limits edit size using ssODNs to ~130 bases (130 + 2x35-base homology arms = 200 bases). To test whether this limit could be surpassed by using overlapping ssODNs, we designed two overlapping ssODNs to generate a 180-base insert at the *gtbp-1* locus. The ssODNs overlapped by ~35 bases and each had a single ~35-homology arm to the DSB. We obtained 63% edits of the correct size (Figure 4A).

#### Large insertions using overlapping PCR fragments

To obtain gene-sized edits, we used overlapping PCR fragments. We designed fragments for *in vivo* assembly of novel GFP fusions, such as GFP fused to a degradation domain (ZF1, 114bp), GFP fused to a membrane localization domain (PH domain, 390bp) and GFP fused to another fluorescent protein (mNeon, 807bp). Fragments overlapped by ~35-bases. We obtained robust editing frequencies up to a combined insert size of 1.6kb (26-52% edits, Figure 4A-D).

When attempting to assemble a larger 2.4kb fusion, however, we obtained significantly lower editing frequencies: 5% edits using two ~1.2kb fragments or three ~0.8kb fragments, and 0% edits using a single 2.4kb fragment (Figure 4E and Supplementary Table 1). The reduced frequency may have been due in part to the lower molarity of each template in the injection mixes (Supplementary Table 1). Using a single GFP template, a ~four-fold reduction in molarity yielded an ~eight-fold reduction in edits (9%, Supplementary Table 1). We conclude that overlapping PCR fragments can be used to assemble repair templates *in vivo*. Best results are obtained when each fragment does not exceed 1kb and the molarity of each fragment is high (~0.5 pmol/μl).

#### GFP-sized insertions using ssODNs to provide homology arms and specify junctions

The use of ssODN to provide homology to the DSB should in principle simplify the construction of templates by eliminating the PCR amplification steps needed to append homology arms to the insert. We first tested this approach by combining a PCR fragment containing GFP with no homology arms and two bridging ssODNs to target the DSB in *gtbp-1*. We obtained between 27 and 53% edits depending on the polarity of the ssODNs (Figure 4F and Supplementary Table 1). As in our earlier experiments using only one bridging ssODN (Figure 3), the most efficient combination involved ssODNs that could pair with the resected template (Supplementary Table 1 and Supplementary Figure 4). We also targeted GFP to the *glh-1* locus, this time using ssODNs designed to insert GFP 6 bases away from the DSB. We obtained 24% GLH-1::GFP edits (Figure 4G, compared to 49% edits using a single PCR template with homology arms (10)). Finally, we also tested whether the bridge ssODN could be used to introduce a short sequence between the PCR template and the chromosomal site. We used an ssODN that included a 66-nucleotides insert coding for a 3Xflag tag to bridge mNeon on the right side of the *gtbp-1* DSB. We obtained 44% edits (Figure 4H). We conclude that ssODNs can be used to target PCR fragments to specific loci and to specify junctional sequences at the insertion point.

#### Gene replacement

We showed previously that it is possible to replace one ORF with a non-homologous ORF using a linear template to repair a deletion generated by two Cas9-induced DSBs (10). The linear template contained the new ORF flanked by homology arms that flank the two DSBs. To test whether bridging ssODNs can be used in this context as well, we replaced the *gtbp-1* ORF with a non-homologous sequence (from the *meg-3* locus) using two ssODNs to provide the homology arms to two Cas9-induced DSBs at either end of the *gtbp-1* ORF. The 3’ ssODN also contained an epitope tag. We obtained 53% edits of the correct predicted size (Figure 4I). We conclude that bridging ssODNs can also be used to delete and replace an ORF in one step.

Next we tested whether this approach could also be used to replace an ORF with a mutated version of the same ORF (so as to introduce multiple mutations at once). When attempting this at the *meg-3* locus, we found that template switching between the recoded template and the locus around each DSB prevented the insertion of all the mutations (data not shown). We therefore developed a two-step strategy. We first deleted the *meg-3* locus using two Cas9-induced DSBs and a bridging ssODN designed to insert a new Cas9 recognition sequence at the junction. We then used two overlapping PCR fragments to reinsert 1.8kb of the recoded *meg-3* ORF at the engineered site. We obtained 6% edits (Figure 4J). The low edit frequency may have been due to the large insert size, the low molarity of each template (0.3pmol//μl), and/or possibly the low efficiency of the engineered Cas9 site.

### Sequencing of edits suggest a precise repair process

We sequenced 76 edits from 21 independent experiments for a total of ~50,000 bases, including 236 novel DNA junctions. We identified 4 mutations in three edits (Supplementary Table S1), including an 8-base indel near the 3’ end of GFP and 3 substitutions. Of the three substitutions, only one mapped to a junction (1 mutation in 236 junctions sequenced). We conclude that the majority (>95%) of edits we report here are precise. In yeast, synthesis-dependent strand annealing is associated with increased mutation rate due to frequent dissociation of the replicating strand from the template (13). Since many edits were identified based on GFP expression, many edits with deletions would not have been recovered. In fact, this possibility could explain why longer inserts tended to yield lower edit frequencies, since longer replication tracks are predicted to experience more dissociation events. Consistent with this possibility, when screening for gene replacement edits by PCR (Figure 4I), we observed at least one incorrectly-sized edit. Sequencing revealed that the edit had precise junctions and an internal 232bp deletion (data not shown).

## DISCUSSION

In this study, we demonstrate that short homologies (~35 bases) are sufficient to support recombination between PCR fragments, ssODNs and Cas9-induced DSBs. Based on these findings, we developed new strategies for genome editing in *C. elegans* using recombineering. We discuss possible mechanisms of recombineering, as well as advantages and limitations for genome editing.

### Repair of Cas9-induced DSBs proceeds by a mechanism involving templated DNA synthesis and template switching

Our results using ssODNs indicate that repair of Cas9-induced DSBs is a polarity-sensitive process that requires annealing of the ssODN to the 3’ strand on one side of the DSB to initiate DNA synthesis, as has been shown in yeast (14). We suggest that PCR templates also participate in a repair process involving synthesis-dependent strand annealing (SDSA). When using PCR templates, two lines of evidence indicate that the initial annealing step is most efficient when involving sequences directly flanking the DSB and sequences at the ends of the PCR templates. First, PCR templates with homology arms corresponding to sequences that directly flank the DSB perform better than templates with homology arms corresponding to sequences located ~30 bases away. Second, extending the homology arms on PCR templates beyond 35 bases reduces editing efficiency, whether the added sequences are homologous or non-homologous to sequences surrounding the DSB. One possibility is that DSBs and PCR fragments are resected by a short-range mechanism that liberates less than ~50 bases of single-stranded DNA. Such a short-range mechanism would be in contrast to resection of DSBs in yeast, which has been proposed to extend over thousands of bases (14). Alternatively, DNA ends could be made available for pairing by another short-range mechanism not involving resection. Homologies internal to PCR templates can also participate in the recombination process (see template switching below), but are not sufficient to support high efficiency gene conversion in templates with only internal homologies (Figure 1), perhaps because of the greater difficulty of invading a paired duplex.

After the initial annealing event and onset of DNA synthesis, our observations indicate that the elongating strand is able to withdraw from the first template and anneal to a homologous sequence in a second template to continue DNA synthesis. Template switching can occur anywhere along a donor template, can occur between two donor templates or between a donor template and the chromosome, and only requires a short homology track (we observed 100% template switching with a 27-base stretch, Figure 1F and 1G). We suggest that, during recombineering, reiterated rounds of annealing/template switching link multiple DNA molecules until a template with homology to both sides of the resected Cas9-induced DSB is assembled and sealing of the DSB is completed. Recombination between free DNAs co-injected in the germline of *C. elegans* has been documented previously (15) (16), raising the possibility that template assembly could be initiated extra-chromosomally before recombination with the Cas9-induced DSB. Kemp et al., 2007 estimated that only 10% of co-injected DNAs recombine (16). In contrast, we observed gene conversion rates as high as 85% at Cas9-induced DSBs (Figure 1), suggestive of a highly efficient repair mechanism initiated by chromosomal DSBs. One possibility is that Cas9-induced DSBs are repaired by the same SDSA-like mechanism that repairs the majority of *spo-11*-induced DSBs that arise during meiosis (17).

### Application of Cas9-assisted recombineering: *in vivo* assembly of knock-in fusions

Taking advantage of the efficiency of gene conversion and template switching in the *C. elegans* germline, we have developed recombineering strategies for the generation of chromosomal inserts without selection. The strategies follow a simple set of rules for template design. First, templates should have short, terminal overlaps, no greater than 35 bases. Homology arms targeting the chromosomal DSB should also be short (~35 bases), located at the end of the template, and at least one should correspond to sequences directly flanking one side of the DSB. Homologies located internally on templates can also participate in the recombination process, and therefore should be avoided to prevent partial inserts. ssODNs and/or PCR fragments can be used as templates. When using only ssODNs, our results so far indicate that ssODNs function best when sharing the same polarity. ssODNs can also be used in combination with PCR fragments to bridge two PCR fragments, or to bridge PCR fragments to chromosomal DSBs. When used in combination with PCR fragments, both polarity of ssODNs are tolerated, although, when targeting GFP to the *gtbp-1* DSB, we observed a preference for ssODNs that can pair and extend the PCR template.

Recombineering using multiple ssODNs make it possible to create edits that would be too large to fit on a standard commercial ssODNs (>130 bases). We do not yet know what is the maximal insert size that can be assembled using only ssODNs. To make gene-sized edits, we used combination of bridge ssODNs and PCR fragments. This strategy eliminates the PCR step needed to append homology arms to templates. For example, for GFP knock-ins, locus-specific ssODNs are injected along with a universal PCR fragment containing GFP. Recombineering can also be used to assemble novel fusions *in vivo*. For example, immunogenic peptides or localization domains can be combined with GFP to create novel, multi-part fluorescent tags. The additional sequences are provided on bridging ssODNs or on overlapping PCR fragments.

### Limitations of Cas9-assisted recombineering

An important requirement of recombineering is that each template DNA must be injected at a molarity sufficient to support efficient repair. A 0.7kb insert generated 75% edits when injected at 0.5pmol/μl, but only 9% edits when injected at 0.1pmol/μl. Because it is difficult to routinely synthesize large DNAs at high molarity, in our hands this limitation resulted in a workable upper limit for total insert size of ~1.6kb (26% edits at the *gtbp-1* locus and 6% edits at the *meg-3* locus). In the future, it may be possible to increase this upper size limit with more concentrated DNA preparations.

The high efficiency of template switching also imposes certain limitations. First, template switching makes it difficult to create edits at a distance from a Cas9-induced DSB (10–12). This is because sequences that span the DSB will promote template switching between the repair template and the chromosome on both sides of the DSB, and prevent the copying of distal sequences on the template (Figure 1F and 1G). This limitation requires that a suitable Cas9 site (with the required PAM sequence) be identified in the vicinity (<20 bases) of every desired insertion site. As new RNA-guided endonucleases with different PAM specificities become available (18,19) this requirement will become easier to satisfy. Second, template switching also makes it difficult to replace an ORF with a closely related sequence in one step, since any internal stretch of homology will promote template switching and prevent the exchange of distal sequence. In that instance, it is preferable to create the edit in two steps: first, generate a deletion that removes the ORF, and second, re-insert the desired fragment (Figure 4J). Finally, template switching may also select against long insertions, since extended replication tracks are more likely to experience aberrant dissociation events leading to deletions and rearrangements (13). Fortunately, protocols already exist to create long edits or edits at a distance from a Cas9 site. These protocols use plasmid-based templates with long homology arms and selectable markers, which make it possible to recover rare recombinants (20). Unlike the approach we described here, plasmid-based methods require cloning and give rise to edits that contain the selectable marker, which must be removed in second step.

### Future prospects

In conclusion, we have shown that Cas9-induced DSBs in *C. elegans* are repaired by a highly efficient gene conversion mechanism that utilizes linear donor DNAs with short homology arms. The gene conversion process can be harnessed for recombineering and yields gene-size edits at high enough frequencies (5% to 85%) to not require selection. The ability to use bridging ssODNs to target GFP to any locus should streamline pipelines for genome-wide GFP-tagging projects. In addition to exciting technical opportunities, recombineering in *C. elegans* also offers a new platform to study the mechanisms that underlie homology-dependent repair in animals. Our observations suggest that, like yeast (9), *C. elegans* possesses a robust mechanism for template switching during repair-induced DNA synthesis. Consistent with this view, evidence for template switching has been observed in DNA rearrangements isolated from telomerase-deficient mutants (21) and from worms exposed to DNA damaging agents (22). We anticipate that, as in yeast and bacteria (6), recombineering in *C. elegans* will expand the already many opportunities afforded by RNA-guided endonucleases.

## SUPPLEMENTARY DATA

All experiments are detailed in Supplementary Table S1. Schematics of experiments with edit examples are in Supplementary Figure S1. Information on the polarity requirements for donor ssODNs can be found in Supplementary Figure S2 and S3. Sequences of bridging ssODNs used to insert GFP at *gtbp-1* locus are in Supplementary Figure S4. Sequences of repair templates, crRNAs, ssODNs and PCR primers in Supplementary Tables S2/S3/S4, and plasmids and strains in Supplementary Table S5.

## ACKNOWLEDGMENTS

We thank Dominique Rasoloson, Andrew Folkmann and Chih-Yung Lee for Cas9 protein purification and plasmid, Andy Fire and Andrew Folkmann for comments on the manuscript, and WormBase for gene annotations and sequences.

